# Unmasking genetic load: Genomic inbreeding drives growth depression in captive critically endangered Chinese Bahaba (*Bahaba taipingensis*)

**DOI:** 10.64898/2026.09.11.750860

**Authors:** Yuxuan Zhang, Lin Yan, Kuoqiu Yan, Runsheng Li, Jilin Zhang, Wenlong Cai

**Affiliations:** Department of Infectious Diseases and Public Health, Jockey Club College of Veterinary Medicine and Life Sciences, City University of Hong Kong, Kowloon Tong, Hong Kong SAR, China; Guangdong Bluegen Marine Biotechnology Co., Ltd, Huizhou, China; Department of Biomedical Sciences, College of Biomedicine, City University of Hong Kong, Kowloon Tong, Hong Kong SAR, China

**Keywords:** Chinese bahaba, conservation genomics, inbreeding depression, genetic load, GWAS

## Abstract

Chinese bahaba (*Bahaba taipingensis*) is a critically endangered marine fish that has been commercially extinct in the wild. Conservation-oriented captive breeding has become an effective strategy for rescuing such endangered species, but concerns about inbreeding depression often remain. Here, after merely two generations of closed breeding from 18 wild founders, Chinese bahaba exhibited severe growth depression, with body length and weight diverging by more than three-fold. To further investigate the early genomic consequences of captive breeding, we analyzed whole-genome sequences from 348 *F*_2_ individuals together with 19 wild individuals from the same marine region. While genome-wide genetic diversity was overall preserved, closed breeding rapidly partitioned the *F* population into eight distinct genetic lineages. A significant elevation in genomic inbreeding was observed to facilitate the homozygous exposure of recessive deleterious mutations, which was strongly correlated with growth decline. An integrative framework combining GWAS, gene-level burden test, and ROH hotspots was applied to dissect the genetic architecture underlying growth decline, which converged predominantly on skeletal scaffolding and immune regulation. This was evidenced by candidate genes related to skeleton development and homeostasis (e.g. *fam20b*, *c-fos*, *trpv5*, and *il23r*), the GH/IGF axis (e.g. *socs1a*), and immune/inflammatory pathways (e.g. *il23r*, *nlrp7*, *cd163*, *il12rb2*), as well as calcium ion transport pathway. Together, our findings provide novel insights into the genomic consequences and genetic architecture of growth decline under captive conditions in Chinese bahaba, and offer critical guidance for genetic management in captive conservation programs of endangered species.

## 1. Introduction

Human activities and climate change have increasingly fragmented natural habitats and threatened the survival of many species worldwide (Scanes, 2018). As a result, many populations that were once large and widespread have been reduced to small, isolated remnants, facing a risk of extinction (Fahrig, 1997). In response to this growing crisis, captive breeding programs, often coupled with reintroduction, have become important conservation strategies for rescuing endangered species and rebuilding wild populations (Dobson and Lyles, 2000, Ewen et al., 2012). This approach is particularly prevalent in aquatic species conservation, where captive propagation and stock enhancement have been widely implemented (Manubens et al., 2020, Roques et al., 2018). However, captive populations are typically founded by only a limited number of wild individuals. Although small, isolated populations may persist for hundreds of generations (Dussex et al., 2021), such demographic conditions, whether in natural or captive settings, inevitably intensify founder effects and genetic drift, resulting in increased genetic differentiation and reduced genetic diversity (González et al., 2020, Forstmeier et al., 2007). More importantly, increasing relatedness expands the extent of genomic regions that are identical by descent (IBD), thereby exposing masked recessive genetic load as realized genetic load (Charlesworth and Willis, 2009). These processes can impair multiple aspects of population performance, including fitness and a range of physiological traits, known as inbreeding depression (Crates et al., 2023). Some studies have shown that significant declines in growth and survival can emerge after as few as two generations of captive breeding (Hasselgren et al., 2024, Li et al., 2024). Such inbreeding-driven deterioration brings challenges to the long-term maintenance of captive stocks and reduces their value for future population reinforcement and reintroduction efforts.

Among fitness-related traits, growth is a key focus in conservation captive breeding and release programs, especially for fishes and other aquatic species, as it is closely associated with individual survival and physiological performance (Primo et al., 2021). Biologically, growth is a complex polygenic trait. In fish, the growth hormone/insulin-like growth factor (GH/IGF) axis serves as a central regulatory system governing both muscle and skeletal development (Fuentes et al., 2013, Hildahl et al., 2008). Muscle growth is further modulated by myogenic regulatory factors (MRFs) and genes involved in energy metabolism, whereas coordinated skeletal development is regulated by the GH/IGF system in together with skeleton-related factors (Johnston et al., 2011, Vieira et al., 2013, Lavajoo et al., 2020). However, the genetic consequences of small, closed breeding populations can readily disrupt these complex physiological regulatory networks. For example, substantial declines in growth associated with high levels of inbreeding have been reported in captive aquatic species such as turbot and Chinese shrimp (Yang et al., 2021, Luo et al., 2014). Moreover, although sensitivity to inbreeding varies among species, growth in aquatic species is considered to be more strongly affected by inbreeding than in many other taxa (Moss et al., 2008). Nevertheless, the detailed genetic mechanisms and population-genetic processes underlying growth variation in small, closed populations remain incompletely understood.

In small, isolated populations that have experienced inbreeding, dissecting how inbreeding and deleterious mutations shape complex phenotypes presents methodological challenges. Currently, genome-wide association studies (GWAS) remain the most widely used approach for elucidating complex growth traits of aquatic species (Omeka et al., 2022, Yoshida and Yáñez, 2021, Tai et al., 2025), owing to their power to detect common polygenic loci of small effect, although the proportion of phenotypic variation explained by the identified signals is often limited (Chaivichoo et al., 2023). For large-effect deleterious mutations that have been randomly enriched by founder effects and unmasked when inbreeding generates runs of homozygosity (ROH), GWAS lacks the statistical power to effectively detect these variants, as they are often rare and poorly captured in standard association models (Szustakowski et al., 2021). In addition, broad genomic regions often contribute collectively to complex traits like growth. Since GWAS is primarily designed to detect discrete variant sites and most associated variants are located in non-coding regions, it struggles to directly map these signals to genes or functional regions underlying trait variation (Maurano et al., 2012, Spence et al., 2026). These limitations highlight the need for a more integrative genomic perspective. In studies of complex traits, such as longevity in humans (Ying et al., 2024) and survival rate in Soay sheep (Stoffel et al., 2021), combining GWAS with assessments of inbreeding and genetic load, such as ROH hotspot analysis and gene-based burden tests, has emerged as a promising approach. Such an integrative framework may help overcome the limitations of individual methods and provide a more robust method for comprehensively resolving the genetic architecture driving growth decline in small close populations.

Chinese bahaba (*Bahaba taipingensis*), a member of the Sciaenidae family, is a critically endangered marine fish endemic to south China, primarily distributed from the Pearl River Estuary to the Yangtze River Estuary (Sadovy and Cheung, 2003). As one of the largest sciaenid fish, its adult individuals can reach around 2 meters in length and weigh up to 100 kilograms (Trewavas, 1977). Historically, it was a highly prized commercial species due to the medicinal value of its swim bladder (Wang et al., 2009). However, decades of overfishing led to a significant decline in wild populations, and by 1997, it was considered commercially extinct (Sadovy and Cheung, 2003). This species was later classified as critically endangered by the International Union for Conservation of Nature (IUCN) Red List in 2006 (Liu, 2020). China listed it as Grade I state protected species in 2021 and launched artificial breeding programs (National Forestry and Grassland Administration, 2021). However, the limited availability of wild broodstock suggests that the captive populations were established from only a few individuals, which raises concerns about genetic health and sustainable development of these populations.

In this study, we investigated the genetic consequences of conservation captive breeding in Chinese bahaba, established from a limited number of wild founders. We focused on 348 *F*_2_ individuals descended from only 18 wild founders. As a small population that had undergone two generations of closed breeding, the *F*_2_ generation is a critical stage at which pronounced divergence in growth phenotypes has been evident. By comparing this *F*_2_ population with previously reported wild populations, we evaluated genome-wide levels of inbreeding and genetic load in the captive *F*_2_ population. Combined with phenotypic data, the effects of inbreeding and genetic load on individual growth were further quantified. In addition, by integrating multiple analytical approaches, including deleterious mutation burden test, GWAS, and ROH dynamics, we identified the key genes and biological pathways underlying growth differentiation. Overall, our study reveals the genomic dynamics of Chinese bahaba under small-population captive breeding and advances our understanding of the genetic basis of growth differentiation. These findings will help optimize captive breeding strategies for Chinese bahaba while also providing broadly applicable insights for the conservation of other endangered species.

## 2. Methods

### 2.1 Sample collection and phenotyping

The fish used in this study were all sexually immature *F*_2_ offspring from the conservation captive breeding programs at Huangjing aquaculture Co. Ltd (Huizhou, China) (license No.: 2024YL0028). These individuals were generated through two generations of random mating from 18 wild founders (8 females and 10 males; weight: 10-12 kg; body length: 90-100 cm) originating from the Pearl River Delta waters, with all fertilized eggs collected on the same day. The hatched *F*_2_ juveniles were raised in a concrete tank measuring 6 meters in diameter and 1.5 meters in depth, at an approximate biomass of 3 kg/m³. The tank was supplied with a natural water source, and no exogenous hormone treatments were applied to any of the fish. The fish were fed manually three times daily, and water conditions were maintained at temperature 24-28.7, 30-33 ppt salinity, and pH 7.2-8.0. After approximately one year, a total of 350 healthy *F*_2_ individuals were randomly selected for this study. Following anesthesia with MS-222 (50 mg/L), buffered with sodium bicarbonate at twice the MS-222 weight, caudal fin samples were carefully harvested using sterilized scissors from those fish and then immediately transferred to –80°C for storage until DNA extraction. In addition, to compare the captive population with its wild counterpart, we included 19 wild Chinese bahaba individuals from the same marine region for which fin-derived sequencing data had previously been published, with an average sequencing depth of 32 × (Cui et al., 2024).

For 350 *F*_2_ individuals, we measured five important growth-related traits, including body length (BL), body depth (BD), total length (TL), head length (HL), and weight (WT) (Fig. S1). For these highly correlated traits, we applied a dimensionality reduction approach to obtain an integrated measure of individual growth performance for subsequent analysis, thereby preserving the major features of the data while avoiding redundant results. After Z-score standardization, principal component analysis (PCA) was performed on the five phenotypic variables with the prcomp function from the stats package v4.3.1 in R, and the first principal component (PC1) was used as a composite growth index because it explained 98.2% of the total phenotypic variation in growth-related traits.

### 2.2 DNA extraction and sequencing

Genomic DNA was extracted from the caudal fin tissues of all collected fish using the sodium dodecyl sulfate (SDS) method (Wang et al., 2020). The quality of the extracted DNA was subsequently assessed. The NanoDrop spectrophotometer (NanoDrop Technologies, Wilmington, DE) and Qubit 3.0 Fluorometer (Life Technologies, Carlsbad, CA, USA) were used to measure DNA concentration and purity, and 1% agarose gel electrophoresis was performed to evaluate DNA integrity. High-quality DNA samples were then sent to Benagen Co., Ltd (Wuhan, China) for library preparation and sequencing. Briefly, DNA libraries with an insert size of 200-400 bp were constructed using the Hieff NGS® OnePot Pro DNA Library Prep Kit V4 following the manufacturer’s instructions. Finally, 150 bp paired-end sequencing was performed on the DNBSEQ-T7 platform (BGI Inc., Shenzhen, China). A total of 5.51 Tb of sequencing data was generated, with an average of 15.74 Gb per sample and an mean sequencing depth of 23 ×.

### 2.3 SNP calling

The raw sequencing reads were first subjected to quality control using fastp v0.23.4 (Chen et al., 2018) to remove adapters, low-quality reads, and ambiguous bases. Clean reads were then aligned to the Chinese bahaba reference genome (CNGB accession No: CNA0072302) using the BWA-MEM algorithm v0.7.17 (Li and Durbin, 2009). Subsequently, duplicate reads in the alignments were identified and removed by employing the MarkDuplicates module in Picard v3.3.0 (https://github.com/broadinstitute/picard). SNP calling was performed using the mpileup function in BCFtools v1.22 (Danecek et al., 2021).

Initial SNP filtering was performed at the sample level. Raw variants were filtered using BCFtools v1.22 to retain only biallelic SNPs with a Phred quality score > 30, mapping quality value > 30, and no indels within 5 bp of the focal site. SNPs with sequencing depths lower than one-third of the average or higher than three times the average, were further removed using VCFtools v0.1.17 (Danecek et al., 2011) to discard sites with abnormally low or high coverage. Following this initial filtering, two captive individuals were excluded from downstream analysis because their genotyping rate was below 0.8. Population-level SNP filtering was performed based on missing rate and minor allele frequency (MAF) using VCFtools v0.1.17, with thresholds tailored to specific downstream analysis. For analysis involving both the wild and *F*_2_ populations, including population structure inference and population genetic statistics, SNPs with a missing rate > 0.1 or a MAF < 0.02 were excluded, resulting in a total of 1,643,551 high-quality SNPs. A relatively lenient MAF threshold was adopted to avoid excluding variants enriched specifically in the wild population. For ROH detection and mutational burden analysis, only SNPs with a missing rate > 0.1 were removed, yielding a total of 1,892,989 SNPs across the wild and *F*_2_ populations. MAF filtering was not applied in these analyses in order to retain rare variants. For analysis focused on the *F*_2_ population, such as GWAS, we retained SNPs with a missing rate < 0.1 and MAF > 0.05, resulting in 1,535,072 high-quality SNPs.

### 2.4 Population genetics statistics and structure

To explore the genetic consequences of early captive breeding in Chinese bahaba, we compared the population genetic characteristics between the wild and *F*_2_ populations. A subsampling approach was applied in the *F*_2_ population to both nucleotide diversity (π) and linkage disequilibrium (LD) decay analysis to control for statistical biases caused by unequal sample sizes. A random subset of *F*_2_ individuals equal to the wild sample size (n = 19) was randomly drawn across 100 iterations. Nucleotide diversity (π) was estimated for each population in non-overlapping 20-kb windows using Pixy v2.0.0 (Korunes and Samuk, 2021). PopLDdecay v3.43 (Zhang et al., 2018) was employed to calculate average LD correlation coefficients (r^2^) within a distance of 20 kb. Mean values of π and r^2^ across 100 iterations were calculated, together with 95% empirical confidence intervals.

For population structure analysis, the 1,643,551 high-quality SNPs were further pruned for linkage disequilibrium with PLINK v2.0.0 (Purcell et al., 2007) using the parameter “––indep-pairwise 50 5 0.2”. Principal component analysis (PCA) was performed using PLINK v2.0.0, and the first four principal components were visualized with the ggplot2 package v3.5.2 (https://ggplot2.tidyverse.org/) in R to assess population stratification. To further characterize the genetic relationships among individuals, we constructed a standardized genetic relatedness matrix (GRM) using GEMMA v0.98.5 (Zhou and Stephens, 2012) with the-gk 2 option. The GRM was visualized as a heatmap using the pheatmap package v1.0.12 (https://CRAN.R-project.org/package=pheatmap) in R, and hierarchical clustering was conducted using the UPGMA (Unweighted Pair Group Method with Arithmetic Mean) method. Subpopulations within the *F*_2_ population were defined by integrating the results from PCA and the GRM. Differences in growth traits among subgroups were evaluated using the Wilcox test.

### 2.5 ROH calling and genomic inbreeding

Runs of homozygosity (ROHs) were identified in both the wild and captive populations using the ––homozyg function in PLINK v2.0.0. The minimum ROH length was set to 100 kb (––homozyg-kb 100), as this threshold has been widely used in fish, mammals, and humans (Chen et al., 2025, Khan et al., 2021, Kirin et al., 2010). Also, this distance substantially exceeded the maximum extent of LD decay in the study populations (Supplementary Fig. S4b). Shorter thresholds would likely capture background homozygosity and therefore reduce the contrast in ROH burden among individuals. We also used the parameters of “––homozyg-snp 50 ––homozyg-density 50 ––homozyg-gap 500 ––homozyg-het 2 ––homozyg-window-snp 50 ––homozyg-window-het 2 ––homozyg-window-missing 5”. The identified ROHs were further classified into different length categories according to their physical length, including 0.1–0.5 Mb, 0.5–1 Mb, 1–2 Mb, 2–5 Mb, and > 5 Mb. The inbreeding coefficient for each individual was estimated as *F*_ROH_, calculated by dividing the total length of ROHs by the total genome length. To assess the effects of genomic inbreeding on growth in the *F*_2_ population, linear mixed models were fitted between *F*_ROH_ and the composite growth index, as well as each of the five measured growth-related traits, with subpopulation identity as a random effect.

### 2.6 Mutation load

The predicted effects of SNPs on protein function were further annotated. Two related sciaenid species, *Larimichthys crocea* and *Miichthys miiuy*, were used as outgroups for ancestral state inference. The genomes of both outgroups were aligned to the Chinese bahaba reference genome using minimap2 v2.30-r1287 (Li, 2018) with the “asm20” parameter. Conserved sites and variants relative to Chinese bahaba were then identified from the alignments using paftools.js from minimap2 v2.30-r1287. For each biallelic SNP in Chinese bahaba, the allele shared by both outgroup species was defined as the ancestral allele, and the alternative allele was defined as the derived allele. Sites at which the two outgroups carried different alleles were excluded from downstream analyses. Polarized SNPs were then annotated using SnpEff v5.3a (Cingolani et al., 2012) and classified into four functional impact categories: high, moderate, low, and modifier. High-impact variants are defined as those predicted to severely disrupt protein function, such as stop-gain or start-loss mutations. Moderate-impact variants alter amino acid sequences and may affect protein functions, such as missense mutations. Low-impact variants were generally not expected to alter protein functions because they do not change amino acid sequences, such as synonymous mutations. Modifier variants are typically located in non-coding regions and are therefore predicted to have no direct effect on protein functions. High– and moderate-impact mutations were defined as deleterious mutations used for subsequent analysis.

To evaluate whether inbreeding in the *F*_2_ population increased the exposure of recessive variation, we quantified the enrichment of homozygous deleterious mutations within ROHs in both the wild and *F*_2_ populations. The enrichment score was calculated as the proportion of homozygous variants located within ROHs divided by the proportion of the genome covered by ROHs. We further assessed the relationship between the number of homozygous mutations and the degree of inbreeding (*F*_ROH_) in the *F*_2_ population using a linear mix model, with subpopulation identity included as a random effect. In addition, to assess the phenotypic effects of homozygous deleterious mutations, we employed a similar linear mix model to regress growth index and five growth traits on mutation counts in each category, analyzing homozygous and heterozygous variant sets separately.

### 2.7 ROH hotspot and selection signature

Genomic regions potentially associated with inbreeding depression were identified by calculating ROH frequency across the genome in non-overlapping 20-kb windows separately for the fastest– and slowest-growing subpopulations. The difference in ROH frequency between the two groups was then computed. To further investigate whether these ROH hotspot regions also showed concurrent shifts in Tajima’s D, we calculated Tajima’s D for the fastest– and slowest-growing subpopulations in 20-kb windows using VCFtools v0.1.17 and then calculated their window-wise difference. We selected the top 5% of ROH hotspot windows showing the largest excess of ROH frequency in the slow-growing subpopulation and intersected them with the top 1% of windows showing the lowest values of ΔTajima’s D. Genes overlapping these intersecting regions were subsequently subjected to GO enrichment analysis using the clusterProfiler package v4.10.1 (Yu et al., 2012) in R.

### 2.8 Burden test

To evaluate the cumulative effects of deleterious mutation exposure at the gene level, we conducted a burden test using the SKAT package v2.2.5 (https://CRAN.R-project.org/package=SKAT) in R. The analysis was restricted to homozygous mutations annotated as having high or moderate impact, and the growth index was used as the response trait. To ensure statistical robustness, only genes harboring at least two such homozygous deleterious variants across the population were included in the analysis. Following these filtering criteria, a total of 5813 genes were retained for the burden test. To account for potential confounding due to population structure, subpopulation assignment and the first principal component from the genetic PCA were included as covariates in the model. Statistical significance was evaluated using false discovery rate (FDR)-adjusted *p*-values, with a threshold of 0.1.

### 2.9 Genome-wide association study and candidate gene prioritization

Genome-wide association analysis for the composite growth index was conducted in the *F*_2_ population using the 1,535,072 high-quality SNPs. A linear mixed model implemented in GEMMA v0.98.5 was employed:

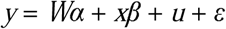

where *y* is a vector containing the trait values, *W* is a matrix of fixed-effect covariates, α is a vector of coefficients corresponding to each covariate in *W*, *x* is a vector of genotype values for a specific genetic marker (SNP), β is the additive effect of that marker, *u* is the vector of random effects accounting for relatedness among individuals, and ε represents random errors. To reduce potential false positives caused by population stratification, the first principal component from PCA was included as fixed-effect covariates in the analysis. Additionally, to account for the effects of genetic relatedness, the relatedness matrix calculated by GEMMA v0.98.5 was incorporated as a random effect. The significance of each SNP was assessed using the Wald test. A genome-wide significance threshold was determined using the Bonferroni correction method (1/N), where N is the total number of SNPs used in the analysis (N = 1,535,072). Given the stringency of the Bonferroni method, a commonly used suggestive threshold of 1 × 10 was adopted (Omeka et al., 2022, Zhu et al., 2023). To identify candidate genes potentially linked to these association signals, we examined the genomic regions within ±10 kb of each suggestive SNP, thereby capturing genes under possible linkage or local regulatory influence.

To further prioritize GWAS signals, we integrated evidence from genomic differentiation between the fastest– and slowest-growing subpopulations. Genome-wide *F*_ST_ was calculated in non-overlapping 20-kb windows using VCFtools v0.1.17, and the top 1% of windows were defined as highly differentiated regions. The top 0.05% most significant GWAS sites were extended by ±10 kb to define candidate association intervals. Genes overlapping both the GWAS intervals and the highly differentiated *F*_ST_ windows were considered prioritized candidate genes.

## 3. Results

### 3.1 Captive-induced genetic stratification and phenotypic segregation

Under closed captive breeding, the 348 *F*_2_ Chinese bahaba individuals analyzed exhibited notable differentiation in the key fitness-related trait of growth. The ranges of head length (HL), body depth (BD), body length (BL), total length (TL), and weight (WT) were 32-117 mm, 31-103 mm, 126-408 mm, 150-478 mm, and 28-855 g, respectively, with mean values of 71 mm, 69 mm, 273 mm, 319 mm, and 297 g (Fig. 1a). The extreme values of these traits differed by more than three-fold. Their coefficients of variation (CVs) ranged from 22.8% to 25.1%, whereas WT showed the greatest variability (CV = 57.7%). All five traits were highly significantly and positively correlated, with Pearson correlation coefficients exceeding 0.96 (Fig. 1a), suggesting that they share a common physiological regulatory basis. Principal component analysis (PCA) on these standardized phenotypes showed that the first principal component (PC1) explained 98.2% of the total growth variation (Supplementary Fig. S2). Therefore, PC1 was employed as a robust composite growth index representing individual growth performance. The growth index showed an approximately normal distribution across the *F*_2_ individuals (Fig. 1a).

**Figure 1.**
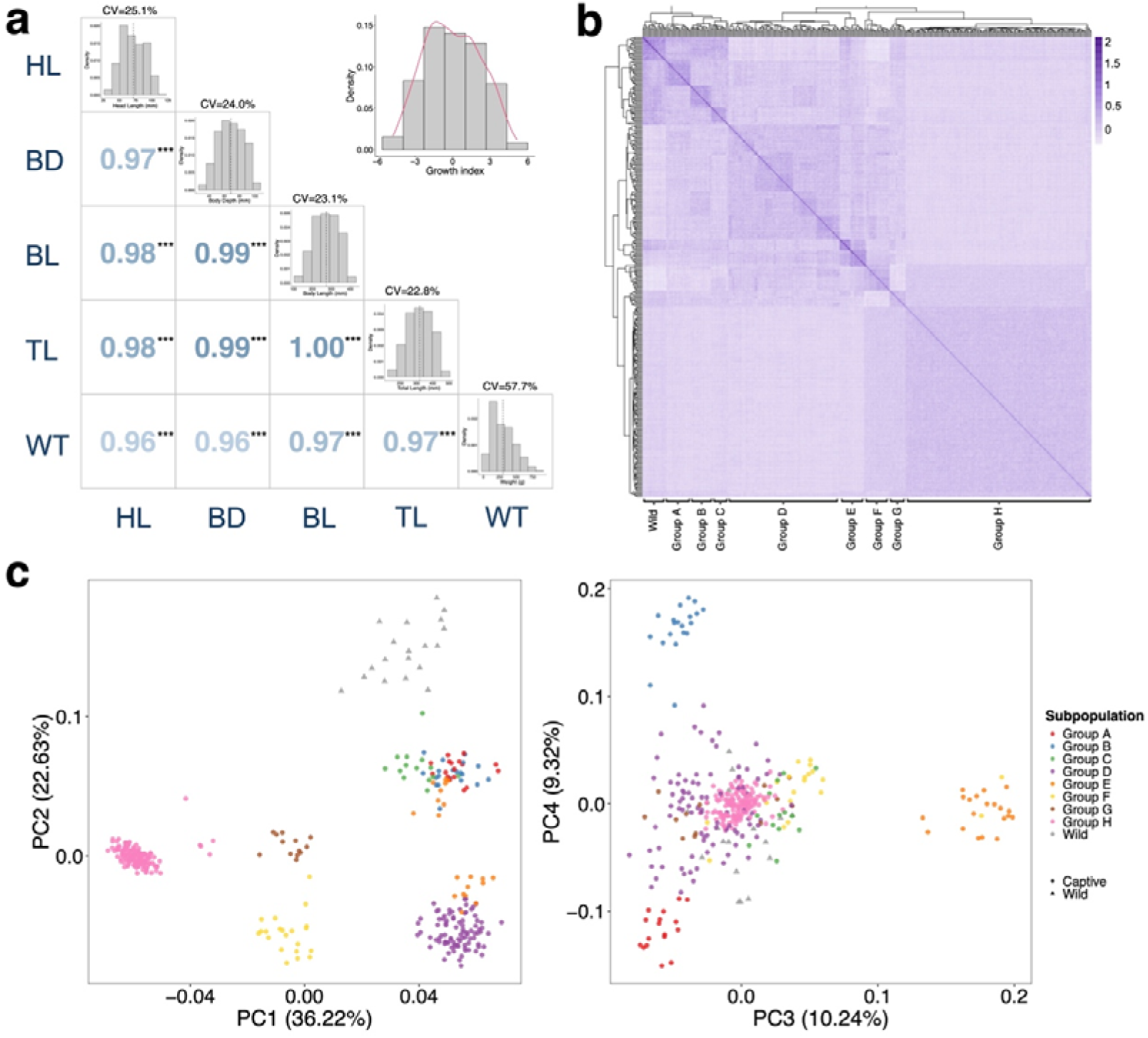
Phenotypic and genetic segregation within the *F*_2_ population. **a.** Phenotypic correlations between growth traits. The lower-left panel reports Pearson correlation coefficients among five growth traits, with significance indicated by asterisks (***, *p*-value < 0.001). Histograms along the diagonal show the distributions of growth traits in the *F*_2_ population, with the dashed line indicating the mean value. The coefficient of variation (CV) for each trait is labeled on each histogram. The histogram in the upper-right corner shows the distribution of the composite growth index in the *F*_2_ population. **b.** GEMMA-derived genetic relatedness matrix (GRM) heatmap with UPGMA clustering tree, displaying clear stratification of the eight *F*_2_ subpopulations. **c.** PCA plots of the first four principal components (PCs), defining eight *F*_2_ subpopulations (Group A-H). In the PCA plot, the ellipses represent the 99% confidence intervals for the wild and *F*_2_ populations, respectively.

Using high-quality SNPs, we reconstructed the genetic relationships and population structure of the *F*_2_ individuals together with the 19 wild individuals. The genetic relatedness matrix (GRM), generated in GEMMA and clustered using the UPGMA method, revealed a highly structured pedigree network and separated the *F*_2_ individuals into eight genetically distinct subpopulations (group A-H) (Fig. 1b), likely corresponding to family lineages descended from the 18 wild founders. Principal component analysis (PCA) showed that the wild individuals exhibited a continuous and dispersed distribution along the first four principal components (Fig. 1c), which reflects a genetically uniform natural population, lacking strong internal substructure. In contrast, the *F*_2_ population was not only separated significantly from the wild individuals but also diverged from the natural pattern, partitioning into several tightly packed subpopulations. Consistent with the GRM results, the first four principal components further supported the presence of eight lineages. Growth phenotypes differed significantly among the eight subpopulations (*p*-value < 0.05), among which group H and group A were identified as the overall fastest– and slowest-growing subpopulations, respectively (Supplementary Fig. S3). It is also noteworthy that unlike the decrease of genetic diversity typically associated with population bottlenecks, the *F*_2_ population retained genome-wide diversity comparable to that of a wild population sampled from the same marine region, whose sample size was similar to that of the wild founders (Supplementary Fig. S4a). In addition, linkage disequilibrium (LD) decay analysis showed that r^2^ in the *F*_2_ population neither decayed more slowly nor remained at a higher level than in the wild population (Supplementary Fig. S4b). These results demonstrate that early-stage captive breeding largely preserved the total pool of natural variation while rapidly segregating it into highly structured lineages.

### 3.2 Elevated genomic inbreeding and ROH dynamics

To trace the genetic consequences of closed breeding, we quantified the accumulation of runs of homozygosity (ROH) for the wild and captive *F*_2_ populations as these regions directly reflect IBD haplotypes derived from inbreeding. Relative to the wild population, the *F*_2_ population showed a notable accumulation of ROHs (Supplementary Fig. S5), indicating a substantial expansion of homozygous genomic regions following only two generations of captive breeding. Classification by ROH length further showed that long ROHs > 1 Mb increased in the *F*_2_ population, with ROHs > 5 Mb even emerging, whereas short ROHs < 500 kb remained predominant, although they were far more abundant than in the wild population (Fig. 2a). This suggests that closed breeding under a limited founder size inevitably increased the burden of identical-by-descent (IBD) segments, but had not yet produced the extensive accumulation of long ROHs typically associated with prolonged inbreeding. Similarly, the proportion of the genome covered by ROHs was significantly higher in the *F*_2_ population than in the wild population, averaging 11.2% and 0.7%, corresponding to *F*_ROH_ = 0.112 and 0.007, respectively (Fig. 2b).

**Figure 2.**
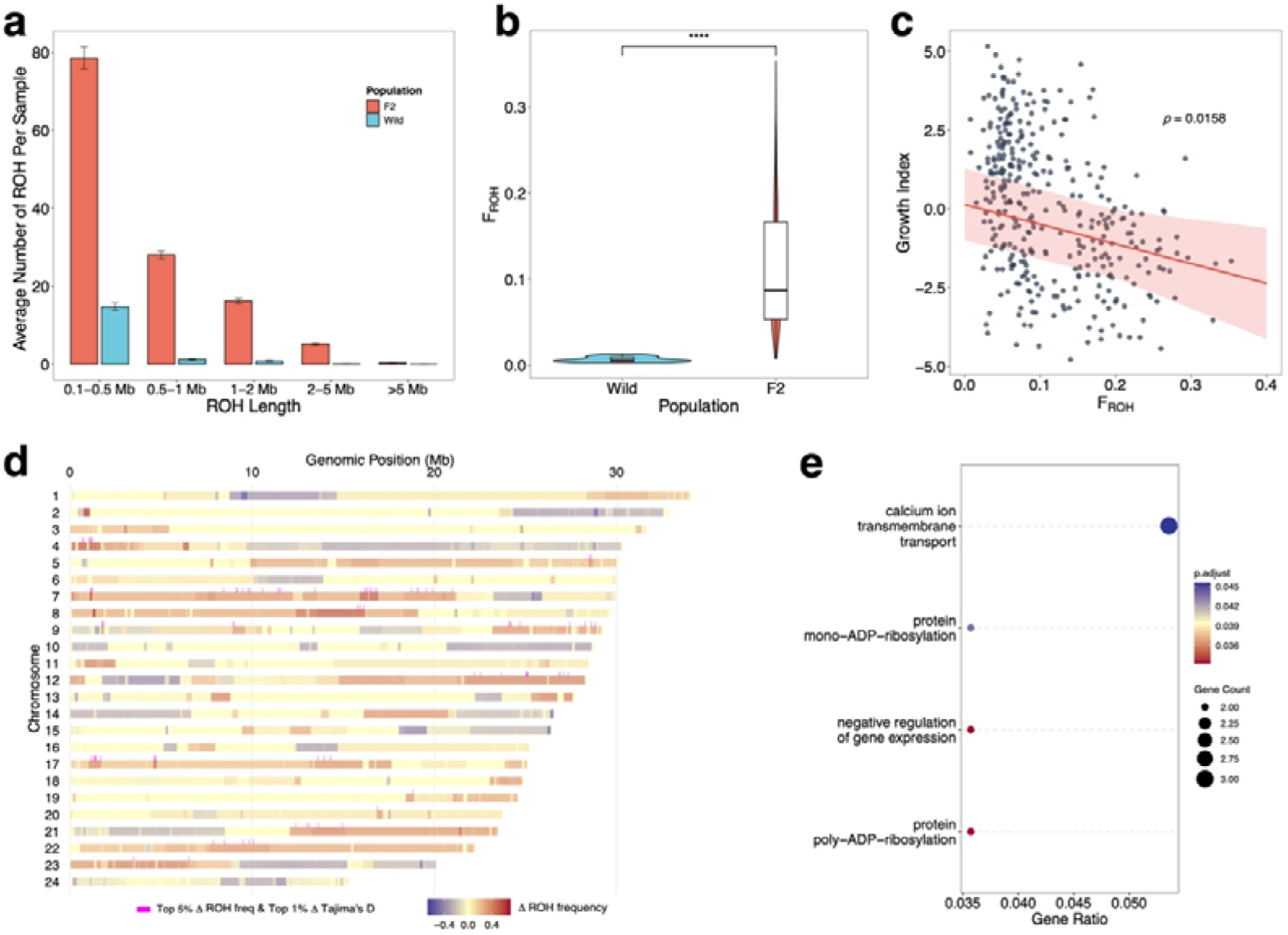
Runs of homozygosity (ROH) accumulation, inbreeding depression, and ROH hotspots. **a.** Average number of ROHs in different ROH length classes in the wild and *F*_2_ populations. **b.** Differences in *F*_ROH_ between the wild and *F*_2_ populations. Significance was assessed using the Wilcoxon test (*, *p*-value < 0.05, **, *p*-value < 0.01, ***, *p*-value < 0.001, ****, *p*-value < 0.0001). **c.** Linear regression showing the negative correlation between individual *F*_ROH_ and growth index. **d.** Differences in ROH frequency in 20-kb windows between the fastest-growing (Group H) and slowest-growing (Group A) subpopulations across the 24 chromosomes. **e.** Gene Ontology (GO) enrichment plot for genes located in the overlapping regions between the top 5% of windows representing slow-growth hotspots and the bottom 1% of windows with the lowest values of ΔTajima’s D_slow-fast_.

Beyond this overall increase, inbreeding levels varied greatly among *F*_2_ individuals, with *F*_ROH_ ranging from 0.008 to 0.353 and a coefficient of variation reaching 63.6% (Fig. 2b). Regression analysis revealed a significant negative relationship between variation in *F*_ROH_ and growth divergence (Fig. 2c and Supplementary Fig. S6), demonstrating genome-wide inbreeding depression. To further characterize such phenotype-specific ROH dynamics, we compared ROH frequencies in non-overlapping 20-kb windows between the fastest– and slowest-growing subpopulations, group H and group A. The genome-wide accumulation of ROHs differed markedly between the two subpopulations and was not uniformly distributed across chromosomes (Fig. 2d). Slow-growth ROH hotspots were mainly located on chromosomes 7, 8, 12, 21, 22 and 23, with the greatest excess of ROH frequency observed in the slowest-growing subpopulation. To further evaluate local genomic shifts associated with these slow-growth ROH hotspots, we compared Tajima’s D in 20-kb windows between the two subpopulations. The top 5% of windows showing slow-growth ROH hotspot signals were intersected with the top 1% of windows showing the lowest values of ΔTajima’s *D*_slow-fast_, overlapping 87 genes (Fig. 2d). These regions therefore represent loci characterized by both homozygosity enrichment and shifts in allele-frequency spectrum between growth-divergent subpopulations. Genes located within these intersecting regions were significantly enriched in pathways related to calcium ion transmembrane transport, a process critical for calcium homeostasis and osteoblastogenesis (Zayzafoon, 2006), as well as protein mono-ADP-ribosylation, negative regulation of gene expression, and poly-ADP-ribosylation (Fig. 2e).

### 3.3 Inbreeding-driven exposure of homozygous deleterious burden represses growth

Strong inbreeding increased the exposure of deleterious mutations. Across the wild and *F*_2_ populations, we successfully annotated 1,604,261 SNPs according to their predicted functional impacts. Among these, variants classified as high or moderate impact accounted for less than 2%, whereas those located in intergenic and intronic regions comprised more than 90% (Supplementary Fig. S7). For high– and moderate-impact mutations, their overall numbers remained comparable between the wild and *F*_2_ populations, but their zygosity changed markedly (Fig. 3a and Supplementary Fig. S8). In the *F*_2_ population, their homozygous mutations increased significantly, accompanied by a significant reduction in heterozygous mutations. In addition, their enrichment scores within ROHs were greater than 1 and significantly exceeded those observed in the wild population (Fig. 3b), indicating that strong inbreeding in the *F*_2_ population facilitated the homozygous accumulation of these variants. These results suggest that two generations of captive breeding may not have generated large numbers of deleterious mutations de novo, but instead promoted the exposure of pre-existing recessive deleterious alleles.

**Figure 3.**
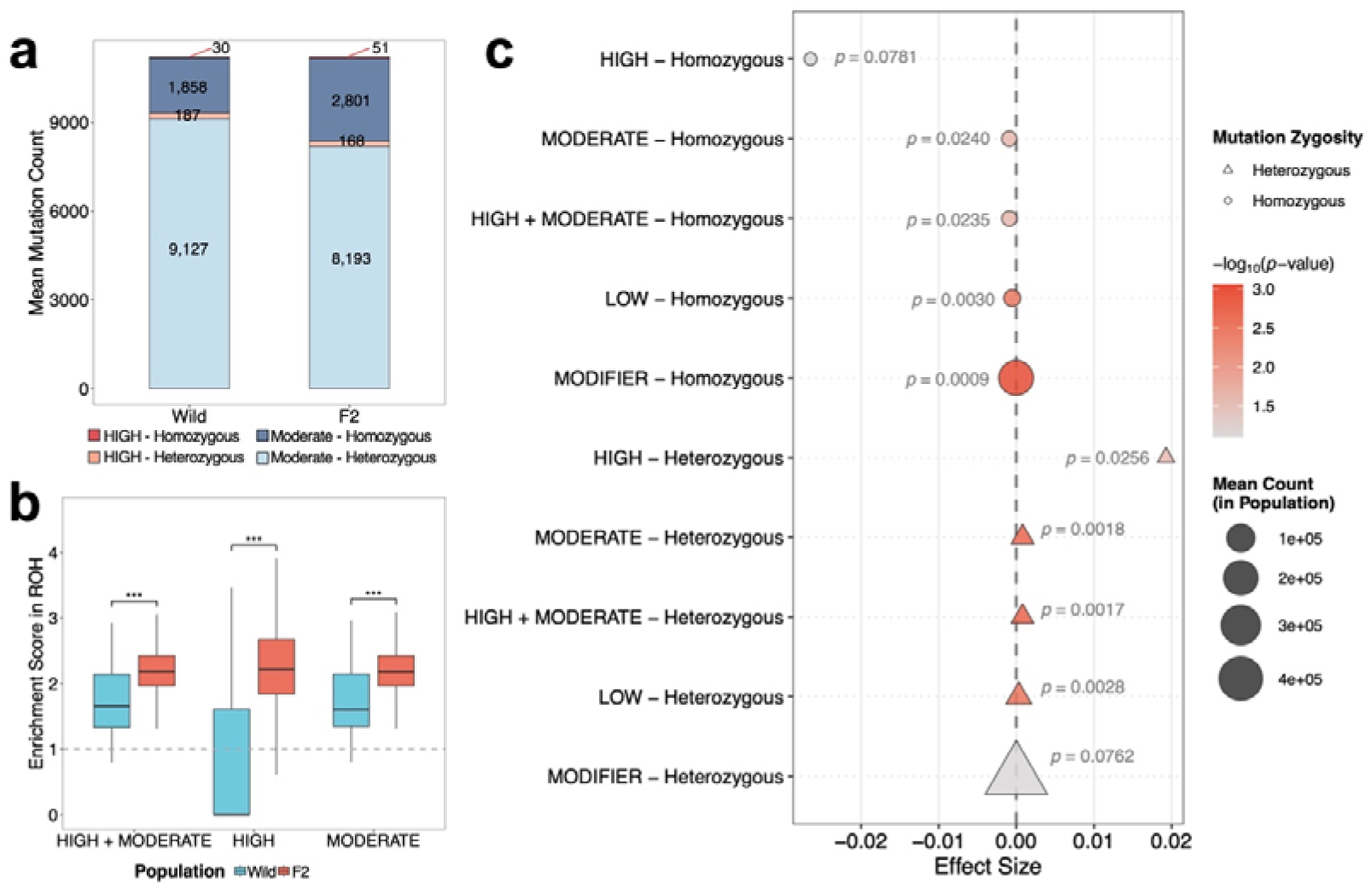
Inbreeding-driven exposure of homozygous deleterious mutations in the *F*_2_ population and association with growth decline. **a.** Stacked bar plot showing the mean numbers of homozygous and heterozygous variants in the four mutation categories annotated by SnpEff in the wild and *F*_2_ populations. **b.** Enrichment scores of homozygous high-impact and moderate-impact mutations located within ROH regions in the wild and *F*_2_ populations. The dashed line at enrichment score = 1 indicates a uniform distribution of homozygous mutations between ROH and non-ROH regions. Significance was assessed with the Wilcoxon test (*, *p*-value < 0.05, **, *p*-value < 0.01, ***, *p*-value < 0.001, ****, *p*-value < 0.0001). **c.** Bubble plot showing regression results in the *F*_2_ population between mutation counts in different mutation categories and individual growth index. Linear mix models were used for all regression analysis.

As expected under inbreeding dynamics, the shift toward increased homozygosity and reduced heterozygosity in the *F*_2_ population extended across all mutation categories. However, the magnitude of these changes varied substantially among *F*_2_ individuals, significantly associated with individual inbreeding levels (Supplementary Fig. S9). To identify the mutation categories contributing to growth decline, we further performed linear regression analysis relating the number of each mutation class to growth index and individual growth traits (Fig. 3c and Supplementary Fig. S10). Importantly, homozygous high– and moderate-impact mutations both exhibited substantial negative effects on growth. Although the scarcity of homozygous high-impact variants across the population limited their individual statistical power (*p*-value > 0.05), pooling deleterious high– and moderate-impact variants improved the statistical significance while maintaining a robust negative association with growth performance. In contrast, while homozygous low-impact and modifier mutations were also negatively associated with growth, their effect sizes were biologically small (approaching zero). Corresponding to the effects observed for homozygotes, heterozygous mutations were positively related to growth performance, indicating that high heterozygosity at functional loci may effectively mask the phenotypic effects of deleterious alleles.

### 3.4 Gene-level burden analysis identifies candidate genes associated with growth decline

Since single-variant association models often lack the ability to capture deleterious alleles that are unmasked collectively across a locus, we conducted a gene-level burden test using SKAT. According to the dominance hypothesis of inbreeding depression, performance decline in captive populations is expected to be driven primarily by the homozygous expression of recessive deleterious mutations (Charlesworth and Willis, 2009). Given their substantial negative effects in our whole-genome regressions, we hypothesized that homozygous high– and moderate-impact variants represent the principal genetic correlates of growth impairment and included them in the burden test. Low-impact and modifier variants were excluded from the analysis, as their little effect sizes would introduce substantial non-functional noise without contributing meaningful biological signals. Growth index was used as the phenotype tested in this analysis. After correction for population structure, the genomic inflation factor (λ) was 1.05 (Supplementary Fig. S11), indicating adequate control of statistic inflation. Following multiple-testing correction (FDR-adjusted *p*-value < 0.1), the burden analysis revealed 18 candidate genes, with a notable concentration on chromosomes 1 and 16 (Fig. 4 and Supplementary Table 1). Among these genes, the majority showed negative effects on the growth index, whereas four genes, including *hnrnpa0*, *ech1*, *socs1a*, and *nbeab*, showed positive effects.

**Figure 4.**
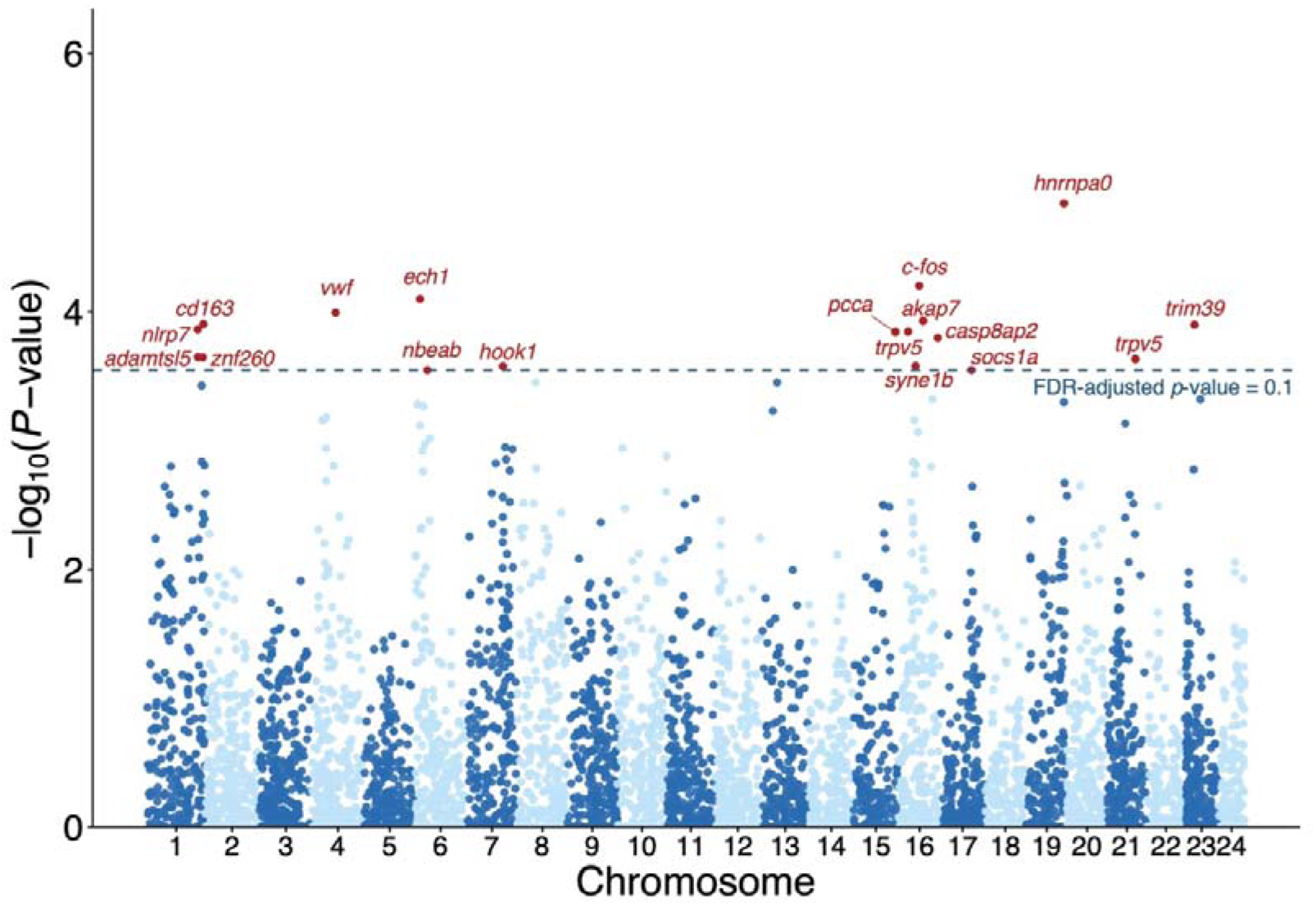
Manhattan plot of the burden test for growth index. The dashed line indicates the significance threshold corresponding to an FDR-adjusted *p*-value < 0.1. Candidate genes exceeding this threshold are highlighted in red.

One noteworthy candidate was *socs1a* positively associated with growth index, which functions as a key negative regulator of the JAK/STAT pathway within the GH/IGF (growth hormone/insulin-like growth factor) axis, and depletion of this gene has been reported to affect post-fertilization growth rate in zebrafish (Dai et al., 2015). In addition to its role in growth regulation, *socs1a* also contributes to the control of systemic autoimmunity (Hanada et al., 2003). Several burden-associated genes were further linked to skeletal development and bone homeostasis. For example, *c-fos* on chromosome 16 encodes a transcription factor that mediates growth factor signaling to promote cell proliferation and differentiation. It has been implicated in osteoblastic differentiation during bone development and remodeling (Ohta et al., 1991, Wagner, 2001). Likewise, two *trpv5* genes on both chromosomes 16 and 21 encode the ion channel protein required for calcium reabsorption and contribute to skeletal homeostasis in humans (Lieben and Carmeliet, 2012). *Adamtsl5* has been detected in skeletal muscle, cartilage, and bone during mouse development (Bader et al., 2012). Other candidate genes in the burden test suggest that metabolic and immune processes may also contribute to growth divergence. Immune– and inflammation-related candidates included *nlrp7*, *cd163*, and *trim39* (Graversen et al., 2002, Chou et al., 2023, Yao et al., 2025), while metabolic candidates included *pcca*, which is involved in amino acid and lipid metabolism (Wongkittichote et al., 2017), and *ech1*, which participates in fatty acid metabolism (Huang et al., 2018). Additionally, *syne1b*, a gene associated with cytoskeletal organization, has been implicated in neuromuscular disease in humans (Baumann et al., 2017).

### 3.5 Joint GWAS and *F*_ST_ analysis prioritizes loci underlying growth variation

To resolve the genomic loci associated with growth variation, we performed a genome-wide association study (GWAS) for growth index on 1,535,072 high-quality SNPs using GEMMA. The heritability estimated by GEMMA was 0.448 ± 0.076. The genomic inflation factors (λ) were 0.98, close to 1 (Supplementary Fig. S12). Applying the suggestive threshold of *p*-value < 1 × 10^-5^, we captured high-confidence signals comprising three SNPs, with two on chromosome 7 and one on chromosome 23 (Fig. 5 and Supplementary Table 2). All three SNPs were located within introns of *anapc4*, *scfd2*, and *intu*, respectively. Gene mapping within their flanking windows (±10 kb) further highlighted *trpv5*, together with *palld* and *ino80b*, as candidate genes (Supplementary Table 2).

**Figure 5.**
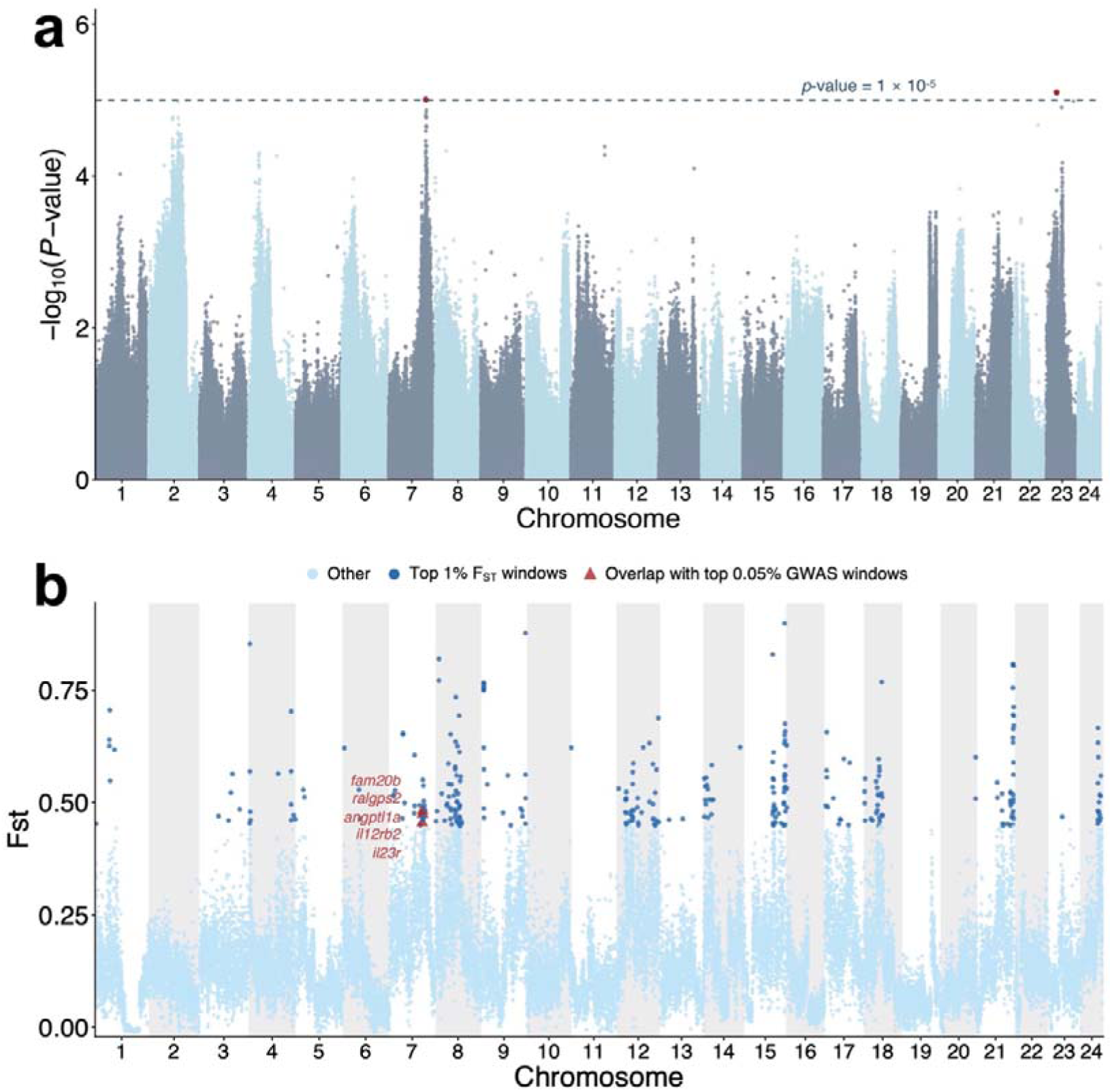
Joint GWAS and *F*_ST_ analysis. **a.** Manhattan plot of GWAS for growth index. The dashed line indicates the suggestive significance threshold of *p*-value < 1×10^−5^. **b.** *F*_ST_ values in 20-kb windows between the fastest– and slowest-growing subpopulations. Dark blue points indicate the top 1% of *F*_ST_ windows. Red triangles indicate the overlap between the ± 10 kb intervals surrounding the top 0.05% GWAS sites and the top 1% *F*_ST_ windows.

Given that a sample size of 348 individuals is limited for complex growth traits, strict thresholding may risk missing polygenic signals. To further capture these signals, we selected the top 0.05% of GWAS sites (n=731 SNPs) and prioritize the key loci by intersecting their ±10 kb windows with the top 1% of *F*_ST_ genomic differentiation windows between the two phenotypic extremes, group H and group A. This integrative filtering successfully identified four robust genomic intervals on chromosomes 7 (Supplementary Table 3), characterized by both strong genetic association and extreme population differentiation. Within these overlapping regions, several key candidates were prioritized, including *fam20b*, *ralgps2*, *angptl1a*, *il12rb2*, and *il23r*. Notably, *fam20b*, *il23r*, and *angptl1a* have been implicated in skeletal development. In zebrafish, *fam20b* deficiency impairs cartilage matrix formation and induces severe skeletal malformations (Eames et al., 2011), while *il23r* has been associated with bone mass during development of mice as an indirect regulator of differentiation of osteoclasts and osteoblasts (Razawy et al., 2021). *Angptl1a*, encoding angiopoietin-like protein 1, has been associated with intermuscular connective tissue and cartilage development (Lai et al., 2007). In addition, *il12rb2* and *il23r*, as interleukin-related receptors, are associated with immune responses (de Paus et al., 2013, Mezghiche et al., 2024).

## 4. Discussion

Captive breeding serves as a vital strategy for preventing the extinction of critically endangered species, but inbreeding depression remains one of the most persistent threats to the long-term success of ex situ conservation programs. Although reduced fitness is widely thought to result from the unmasking of recessive deleterious alleles (Charlesworth and Willis, 2009), the genomic architecture through which inbreeding leads to phenotypic deterioration remains poorly understood. Here, by analyzing 348 *F*_2_ individuals of the critically endangered Chinese bahaba (*Bahaba taipingensis*) derived from a limited wild founder population, we show that remarkable growth decline had already emerged after only two generations of captive breeding. Critically, we observed the homozygous unmasking of pre-existing deleterious alleles by elevated genomic inbreeding, rather than the accumulation of new mutations, which is expected under the dominance hypothesis. Our study provides a valuable opportunity to dissect the genomic basis of early fitness deterioration under captive conditions.

### 4.1 Skeletal scaffold and immune function may form bottleneck limiting growth

Growth in teleosts is predominantly studied in terms of muscle mass accumulation. Here, to overcome the limitations of single-marker analysis and capture the genetic determinants of growth variation, we coupled genome-wide association studies (GWAS) with ROH hotspot detection and gene-level burden tests. Remarkably, these independent analytical methodologies did not converge mainly on muscle-specific pathways, but instead repeatedly highlighted processes related to skeletal scaffold and immune.

It has been suggested that continued body growth also depends on the coordinated development of the skeleton, as it provides the structural scaffold required to support and accommodate muscle growth (Salhotra et al., 2020, Vélez et al., 2018). Nordgarden et al. (2006) found that during the post-smolt growth stage of Atlantic salmon, growth hormone (GH) and insulin-like growth factor (IGF) receptors were upregulated in both muscle and skeletal tissues, indicating coordinated growth and development of them. Our results are consistent with this view. We identified candidate genes related to the GH/IGF pathway, such as *socs1a*, that are involved in both muscle and skeletal growth, and skeletal growth-related functional categories were also represented across multiple candidate loci. Rather than acting within a single pathway, the genes prioritized in burden test and GWAS collectively reveal several interconnected components of skeletal growth, including osteoblastogenesis and osteoclastogenesis (e.g. *c-fos* and *il23r*), calcium transport and homeostasis (e.g. *trpv5*), and cartilage matrix formation and maintenance (e.g. *angptl1a* and *fam20b*). Together with the enrichment of calcium ion transport pathways in slow-growth ROH hotspots, these signals consistently support the view that captive growth depression in Chinese bahaba may result from defects across skeletal scaffold development and homeostasis.

Several candidates identified by both burden testing and GWAS are also functionally associated with immune regulation and inflammatory responses (e.g. *nlrp7*, *cd163*, *trim39*, *il12rb2*, and *il23r*), raising the possibility that variation in immune processes also play a role in growth divergence. This is biologically plausible because immune activity is energetically costly and is often subject to trade-offs with other life activities under conditions of limited energy availability (Wolowczuk et al., 2008, Sheldon and Verhulst, 1996). Studies in humans and plants have suggested that growth and immune function can compete during development (Urlacher et al., 2018, Wang and Wang, 2014). Therefore dysregulation of immune-related pathways may further constrain growth, either directly through inflammatory effects on development or indirectly by diverting resources away from growth-related activities.

### 4.2 Apparent maintenance of genetic diversity in captive conservation

Genetic diversity and inbreeding are widely regarded as key parameters monitored in conservation breeding programs, as they are critical to both genetic health and feasibility of future reintroduction (Frankham et al., 2017). Following population bottlenecks, increased inbreeding and genetic drift often contribute to the loss of genetic diversity, which is closely associated with adaptive potential (Pečnerová et al., 2021). Such reductions in genetic diversity have been widely observed in populations of endangered species, such as aoudad and Grauer’s gorillas (van der Valk et al., 2019, Pizzigalli et al., 2024). However, the paradox regarding genetic diversity has been reported. For example, in isolated populations of kākāpō, genetic diversity was substantially reduced, but deleterious variants remained at a low level due to intensified inbreeding and long-term purging (Dussex et al., 2021). In island fox, populations remained healthy and persist over the long term despite extremely low genetic diversity (Robinson et al., 2016).

In our study, genetic diversity in the captive *F*_2_ population remained comparable to the wild population. This pattern should be interpreted with caution, because wild Chinese bahaba populations have experienced severe demographic contraction in recent decades (Sadovy and Cheung, 2003), and the contemporary wild population may therefore already represent a genetically reduced baseline. More importantly, our results show that with rapid pedigree fragmentation and unmasking of recessive deleterious mutations, substantial genetic changes occurred and already produced differences in breeding performance even before measurable losses of population-wide diversity became evident. This suggests that in the early-generation closed breeding program with limited founders, founder alleles may be primarily redistributed into distinct lineages rather than immediately eliminated from the genetic pool. However, mating within restricted lineages then generated extensive IBD chromosomal regions, driving individual *F*_ROH_ values to as high as 0.35. The resulting increase in homozygosity likely exposed recessive deleterious alleles within ROH blocks, thereby contributing to the growth depression. Therefore, in captive conservation programs, whole-population summary statistics such as genetic diversity alone fail as early warning indicators for population genetic health. Instead, a more comprehensive evaluation incorporating multiple indicators, such as the accumulation of realized genetic load, may supplement the biologically authentic assessment of early-stage captive endangerment.

### 4.3 Inefficient purging of deleterious mutations

Previous studies have debated whether isolated small populations can effectively purge deleterious mutations. Some studies have suggested that inbreeding promotes the expression of deleterious alleles and therefore facilitates their removal (Stuart et al., 2025), whereas others have reported the opposite pattern (Robinson et al., 2016). In our study, homozygous deleterious mutations with high– and moderate-impact annotations were significantly increased in the surviving *F*_2_ population. These mutations were more strongly enriched within ROH, and increased further with rising *F*_ROH_. These patterns suggest that deleterious mutations may not have been efficiently purged, or that some of their effects were too small to be strongly selected against.

Here, the captive population may have a small effective population size as it was founded by only 18 wild individuals and divided into eight lineages. Under this condition, genetic drift may have outweighed purifying selection, leading to an increase in the frequency of deleterious alleles (Hedrick, 1994). Also, under protected aquaculture conditions, artificial feeding and stable environments often reduce survival pressure, which may further limit the efficacy of natural selection (Robert, 2009). In our captive population, homozygous deleterious mutations within skeletal and immune-related genes identified for growth decline, which might otherwise cause incomplete skeletal development or immune abnormalities in the wild, may not be subject to strong purifying selection under captive conditions. Therefore, such retained deleterious variants should be treated with caution in captive populations intended for reintroduction. It is also worth noting that the deleterious mutations analyzed here were inferred from their predicted functional effects on proteins, and such predictions do not always reflect their true biological consequences. Predicted deleterious mutations may even have protective or beneficial effects. For example, in humans, some variants predicted to be deleterious have been associated with disease resistance and longevity (Simon et al., 2022, Freudenberg-Hua et al., 2014), and in livestock species such as cattle, function-disrupting variants in the *mstn* gene increased muscle mass (Grobet et al., 1997). Some mutations that are harmful in one context may also be retained within populations because they contribute to environmental or ecological adaptation in another. This is well supported by the study of Burga et al. (2017), where deleterious developmental mutations underlying skeletal ciliopathies were maintained in flightless Galapagos cormorants to facilitate wing reduction and skeletal adaptation. Similarly, our burden test identified genes carrying homozygous deleterious mutations that showed positive associations with growth, such as *socs1a*. Although *socs1a* is also important for immune homeostasis, deleterious variants it harbours may have been retained due to potential adaptive benefits and relaxed selection pressure.

### 4.4 Limitations and implications for captive genetic management

Our study also has several limitations. First, given that both GWAS and the burden test are susceptible to confounding from population structure, we performed comparisons between phenotypic-extreme subpopulations in ROH hotspots, Tajima’s D, and *F*_ST_, which can reduce false positive signals from such stratification while capturing genomic differences. However, in this recently derived *F*_2_ captive population, Tajima’s D and *F*_ST_ may not reflect selection alone, but also founder effects, genetic drift, and lineage-specific haplotypes. Nevertheless, when combined with other genomic signals, these statistics remain informative for identifying regions associated with growth-related variation. Second, although all captive individuals used in this study were reared under identical conditions, growth is a plastic trait that can be affected by subtle environmental differences and hierarchies, such as feeding competition, which may further influence performance. Third, some deleterious variants annotated here may not always match their true biological effects. Therefore, further functional validation is needed.

Despite these limitations, our findings provide useful guidance for ex situ conservation programs. Genetic management in captivity should consider multiple indicators together, including realized genetic load, especially ROH and the burden of homozygous deleterious mutations, as routine measures of population health. Also, in addition to minimizing relatedness, captive breeding programs should move toward genome-informed mating designs that avoid pairing shared deleterious haplotypes, especially in pathways related to key fitness traits. Founder enrichment and genetic rescue may be needed for long-term viability of captive Chinese bahaba. For reintroduction, it is important to conduct genetic screening before release to help ensure the physiological capacity of released individuals needed for survival and adaptation in the wild and reduce the risks of introducing high-frequency deleterious variants into the wild gene pool.

## 5. Conclusion

Overall, this study demonstrates that closed captive breeding of Chinese bahaba triggered rapid and severe growth depression within merely two generations by unmasking recessive deleterious variants with increased inbreeding levels, even while genome-wide nucleotide diversity remained at an apparently healthy level. Our integrative genomics reveals that this phenotypic divergence may result from variation in genes and pathways related to skeletal scaffolding and immune functions rather than merely muscle growth. These findings highlight the importance of multi-metric genetic monitoring for population health in captive conservation programs, and revealed the necessity of genetic load-aware assisted breeding and genomic screening to mitigate deleterious homozygosity and safeguard individuals intended for wild stock enhancement.

## Ethics statement

All animal procedures were approved by the City University of Hong Kong Animal Care and Use Committee (AN-STA-00001101). We have complied with all relevant ethical regulations for animal use.

## Author contributions

**Yuxuan Zhang:** conceptualization, methodology, formal analysis, investigation, visualization, resources, writing – original draft, writing – review & editing. **Lin Yan:** methodology, data curation, visualization, writing – review & editing. **Kuoqiu Yan:** resources, conceptualization, writing – review & editing. **Runsheng Li:** conceptualization, methodology, writing – review & editing. **Jilin Zhang:** conceptualization, data curation, writing – review & editing. **Wenlong Cai**: conceptualization, methodology, supervision, funding acquisition, writing – review & editing.

## Acknowledgement

We thank the staff at Guangdong Bluegen Marine Biotechnology Co., Ltd for their assistance with sample collection.

