## Supplementary Fig. for "Unmasking genetic load: Genomic inbreeding drives growth depression in captive critically endangered Chinese Bahaba (*Bahaba taipingensis*)"

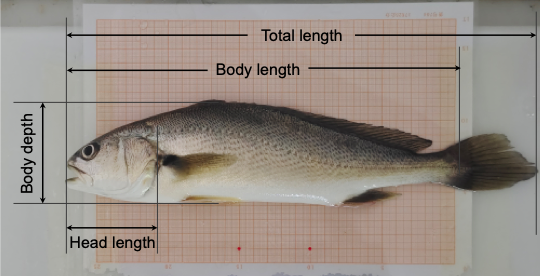

**Figure S1.** Measurement of growth-related traits.

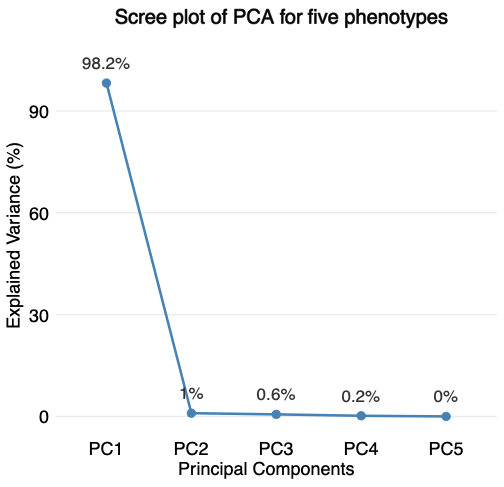

**Figure S2.** Scree plot showing the percentage of variance explained by the first five principal components from the PCA of the five growth traits.

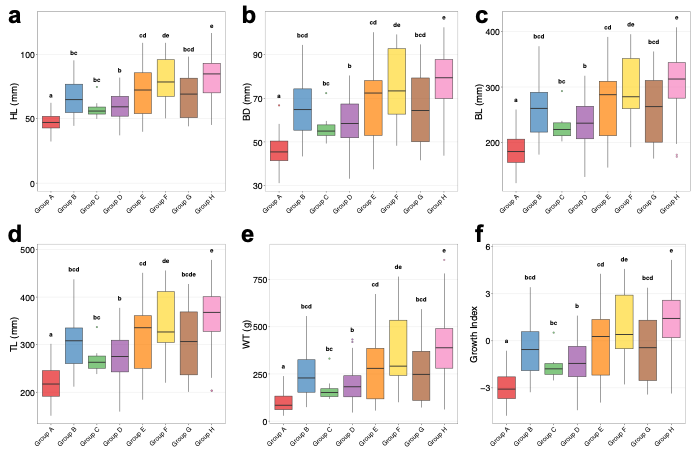

**Figure S3.** Distributions of five key growth-related traits, including head length (**a**), body depth (**b**), body length (**c**), total length (**d**), and weight (**e**), as well as the composite growth index (**f**) across the eight F2 subpopulations. Letters above the boxplots indicate significant differences among subpopulations (Wilcoxon test, *p*-value < 0.05).

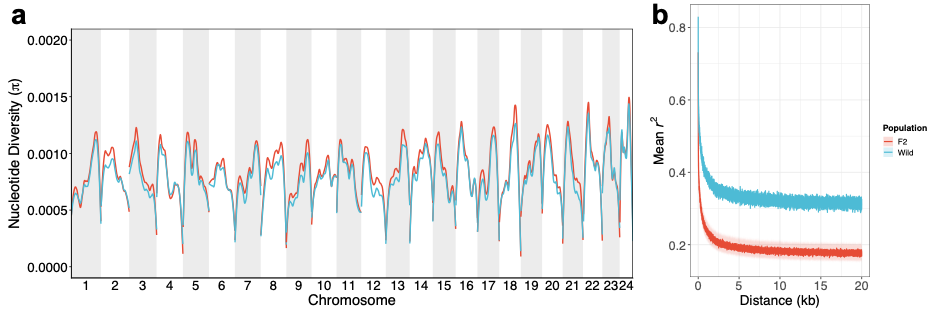

**Figure S4.** Genetic statistics for the wild and F2 population. **a.** Genome-wide genetic diversity (π) estimated in 20-kb windows. For the F2 population, the mean diversity for each window across 100 random resamplings of 19 individuals was calculated. The diversity curves were smoothed using the “loess” function in R. **b.** Linkage disequilibrium (LD) decay curves. The shading around the F2 population curve indicates the 95% confidence interval based on 100 random resamplings of 19 individuals.

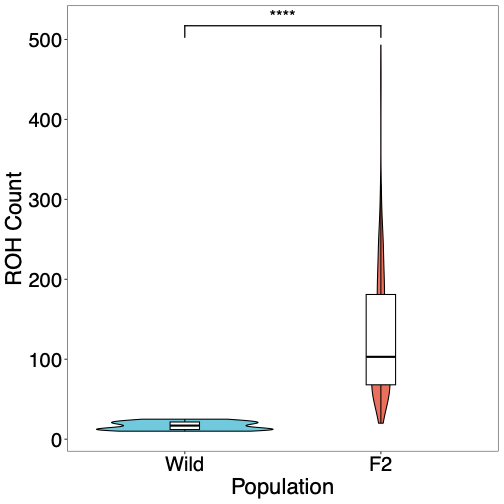

**Figure S5.** Differences in ROH number between the wild and *F*_2_ populations. Significance was assessed using the Wilcoxon test (*, *p*-value < 0.05, **, *p*-value < 0.01, ***, *p*-value < 0,001, ****, *p*-value < 0.0001).

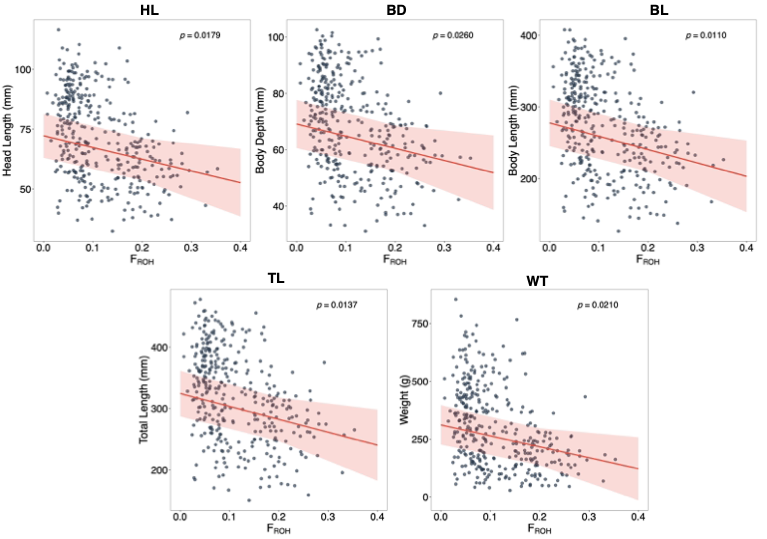

**Figure S6.** Relationships between individual *F*_ROH_ and five growth-related traits in the *F*_2_ population. The shading around the fitted lines represents the 95% confidence intervals.

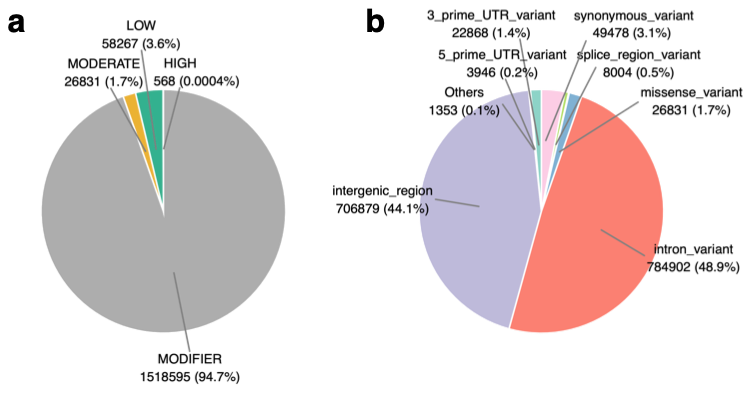

**Figure S7.** Summary of all mutation types annotated by SnpEff in the wild and F2 populations. **a.** Mutations classified into high-, moderate-, low-impact, and modifier categories. **b**. Mutations classified by specific annotation categories.

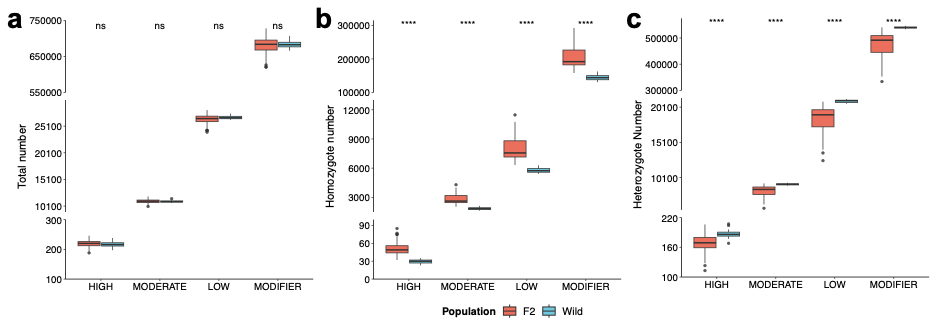

**Figure S8.** Comparison of high-, moderate-, low-, and modifier mutations in their total number (**a**), homozygous (**b**), and heterozygous (**c**) subsets, between the wild and F2 populations. Significance was assessed using the Wilcoxon test (ns, p-value ≥ 0.05, *, *p*-value < 0.05, **, *p*-value < 0.01, ***, *p*-value < 0,001, ****, *p*-value < 0.0001).

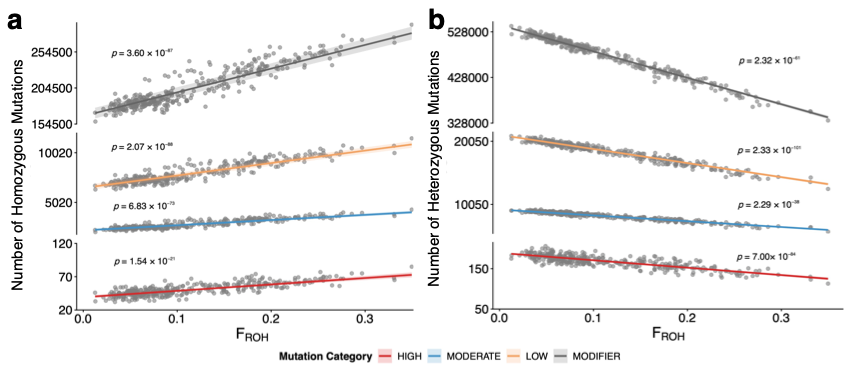

**Figure S9.** Relationships between the number of homozygous **(a)** and heterozygous **(b)** mutations and *F*_ROH_ in the F2 population. The shading around the fitted lines represents the 95% confidence intervals.

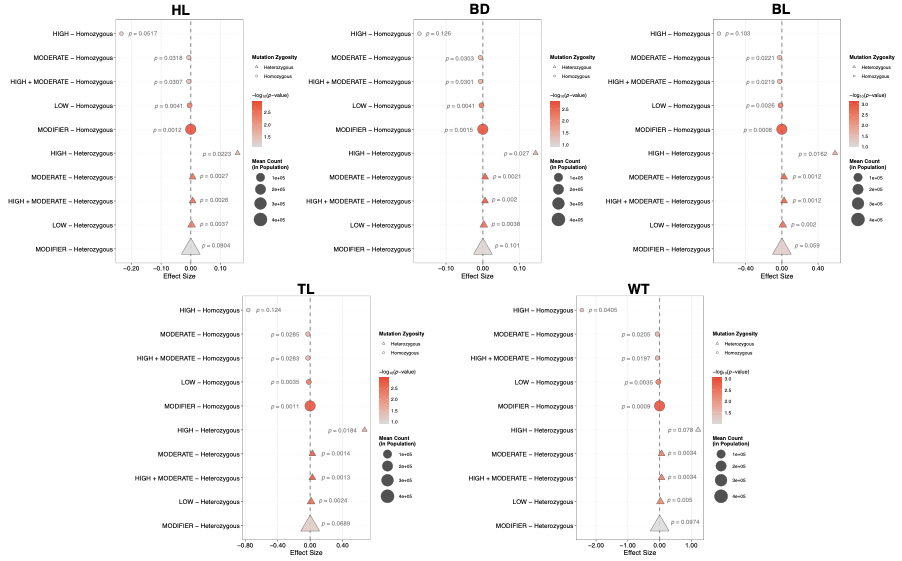

**Figure S10.** Bubble plot showing the regression results for associations between mutation counts in different mutation categories and five growth-related traits in the *F*_2_ population.

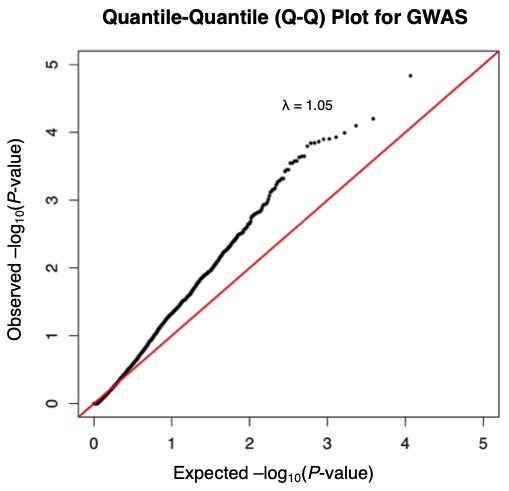

**Figure S11.** Quantile–quantile (QQ) plot of burden test for growth index.

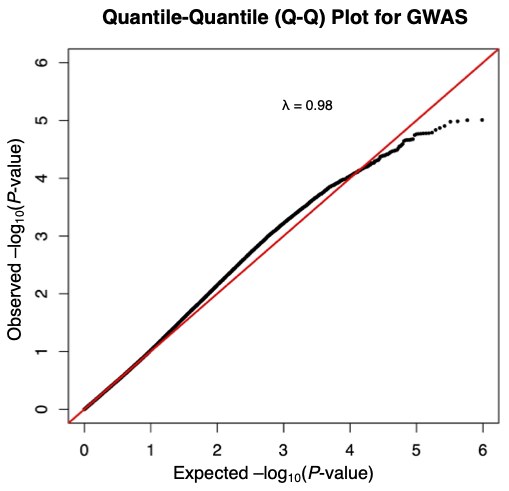

**Figure S12.** Quantile–quantile (QQ) plot of GWAS for growth index.
